# Physiological Characteristics of Viable-but-nonculturable *Vibrio parahaemolyticus* upon Prolonged Exposure to the Refrigerator Temperature

**DOI:** 10.1101/294744

**Authors:** Jae-Hyun Yoon, Sun-Young Lee

## Abstract

Although it has been reported that **v**iable-but-nonculturable (VBNC) cells of *Vibrio parahaemolyticus* can be developed by a prolonged duration of cold-starvation there are restricted cellular characteristics available on understanding the exact mechanisms governing the entry of pathogens into the VBNC state. Therefore, this research was aimed at determining the cellular profile of VBNC cells of *V. parahaemolyticus* upon exposure to the refrigerator temperature. Strains of *V. parahaemolyticus* were incubated in artificial sea water (ASW) microcosms (pH 6) added with different amounts of NaCl at 4°C until these pathogens entered into such a dormant state. At a regular time-interval, both culturability and viability of these bacteria were enumerated, and then cellular profiling were carried out in terms of cellular membrane permeability, enzymatic activity, hydrophobicity, fatty acid composition, and morphological changes after cells of *V. parahaemolyticus* became the VBNC state. Three strains of *V. parahaemolyticus* used in this study showed that VBNC cells retained the strong virulent properties to Vero and CACO-2 cell lines, re-gained the cytotoxicity even after resuscitation, became permeabilized in terms of the outer membrane, showed lower levels of enzymatic (catalase and glutathione-S-transferase) activities, exerted the increasing hydrophobicity, and then exhibited increasing amounts of saturated fatty acids.

**IMPORTANCE:** To the current best knowledge, there are restricted information available on understanding the physiological characterization of viable-but-nonculturable cells. Most previous studies are still making a degree of efforts in discovering the causative effector causing microorganisms to be induced into the VBNC state. Herein, the present study showed that pathogenic *V. parahaemolyticus* can enter into the VBNC state when challenged by a certain environmental stress where higher amounts of NaCl combined with acidic pHs was artificially controlled. Importantly, it was indicated that VBNC *V. parahaemolyticus* maintained peculiarly different physiological characteristics. Furthermore, this study proposed a novel approach on the transient/stepwise conversion of the bacteria into the VBNC state. Specific alternative tools for measuring and controlling the incidence of VBNC pathogens on food are not established until now. In this aspect, results obtained from this study will used to provide an effective insight in determining physiological properties of viable-but-nonculturable *V. parahaemolyticus*.

## INTRODUCTION

*V. parahaemolyticus* has been recognized as one of the major food-borne pathogens commonly found in estuarine environments such as seawater, costal area, and marine sediment (1-2). It has been also reported that human pathogens such as *V. parahaemolyticus*, *Vibrio vulnificus*, and *Vibrio cholerae* can be isolated from a wide variety of raw aquatic products, including clam, mussel, oyster, scallop, and shrimp (3-4). Consumption of marine products contaminated with these pathogens could result in serious clinical symptoms, ranging from acute abdominal pain, vomiting, and nausea to fatal septicemia (5). Especially, it has been determined that cells of *V. parahaemolyticus* are able to enter into a viable-but-nonculturable (VBNC) state in response to a physiological ecology such as higher amounts of NaCl (6) and low temperatures of ≤10°C (7). Until now, more than 65 species of microorganisms, including *V. parahaemolyticus*, *V. vulnificus*, *V. cholerae*, *Campylobacter jejuni*, *Escherichia coli* O157:H7, *Listeria monocytogenes*, *Salmonella enterica* serovar Typhimurium, *Shigella dysenteriae*, and *Staphylococcus aureus* are proven to be induced into the VBNC state upon exposure to environmental stresses (8-13). It should be noted that VBNC bacteria cannot represent the colony-forming capability on agar plates, on which these organisms can grow routinely, thereby indicating that viable-but-nonculturable *V. parahaemolyticus* might not be detected by means of the standardized cultivation-based methods. Although lots of studies have been conducted to find out the exact mechanisms governing the entrance of bacteria into such a dormant state there are still restricted scientific evidences available on understanding physiological characteristics of VBNC organisms. After producing VBNC *V. parahaemolyticus*, these bacteria showed a significantly higher resistance to ethanol shock, as compared with the actively growing cells cultured in tryptic soy broth (14). Above all, it was demonstrated that co-culturing VBNC pathogens such as *V. parahaemolyticus* and *Shig. dysenteriae* with clinical cell lines resulted in reducing a magnitude of viable animal cells, disrupting the cellular substructure undiscriminatingly (13, 15). Thus, it appeared that VBNC pathogens would be associated with the food-borne disease outbreaks, thereby threatening public health concerns. Nevertheless, it might be difficult to construct an effective microbial risk assessment for controlling the pathogenic bacteria due to a significant lack of cellular properties of the VBNC cells. In the present study, cellular properties of viable-but-nonculturable *V. parahaemolyticus* were characterized by measuring cytotoxic effects to human cell lines, the membrane permeabilization with ß-galactosidase and ß-lactamase assays, intracellular leakages of nucleic acid and protein, enzymatic activities of catalase and glutathione-S-transferase (GST), and the cellular hydrophobicity. Meanwhile, cellular fatty acid composition assays were conducted between pure cultures and VBNC cells of *V. parahaemolyticus* for the purpose of revealing the kinetic of mechanisms mediating the conversion of bacterial cells into the VBNC state. Furthermore, not only the ability of *V. parahaemolyticus* strains to be recovered from the VBNC state, but morphological changes before and after the entry into the VBNC state were examined to develop a well-standarized mean for determining the physiological characterization of *V. parahaemolyticus* in the VBNC state.

## RESULTS AND DISCUSSION

### Loss of the culturability of *V. parahaemolyticus* strains

In order to induce cells of *V. parahaemolyticus* into the VBNC state, three strains of *V. parahaemolyticus* ATCC 17082, *V. parahaemolyticus* ATCC 33844, and *V. parahaemolyticus* ATCC 27969 were incubated in ASW microcosms (pH 6) supplemented with different amounts of NaCl at 4°C, respectively. Bacterial populations were enumerated at regular time-intervals by means of the cultivation-based method on agar plates. As shown in Fig. 1, initial densities of *V. parahaemolyticus* ATCC 17082, *V. parahaemolyticus* ATCC 33844, and *V. parahaemolyticus* ATCC 27969 ranged from 6.0 log_10_ CFU/ml to 8.0 log_10_ CFU/ml. Populations of *V. parahaemolyticus* ATCC 17082 exceeded approximately 4.0 log_10_ CFU/ml until incubated in ASW microcosms containing 0.75% NaCl or amended with 5% NaCl at 4°C for 40 days, whereas microbial loads of *V. parahaemolyticus* ATCC 17082 dropped to below the detectable levels (<1.0 log_10_ CFU/ml) at 4°C within 80 days. Especially, loss of the culturability of *V. parahaemolyticus* ATCC 17082 was accelerated more progressively at increasing NaCl concentrations in ASW solutions, showing that cells of *V. parahaemolyticus* ATCC 17082 in ASW microcosms added with 10% and 30% NaCl were induced into the nonculturable state at 4°C within 21 days. There were no differences in the bacterial reductions between three strains of *V. parahaemolyticus* incubated in ASW microcosms containing 0.75% NaCl or supplemented with 5% NaCl. During the cold-starvation challenge, the culturability of these pathogens declined below the detectable levels within 80 days. Within 21 days, cells of *V. parahaemolyticus* ATCC 33844 entered into the nonculturable state in ASW microcosms added with ≥10% NaCl at 4°C. Cells of *V. parahaemolyticus* ATCC 27969 were also converted into the nonculturable state when incubated in either formal ASW microcosms or those amended with 5%, 10%, and 30% NaCl at 4°C for 7, 21, 80, and 80 days, respectively. These results indicated that higher degrees of NaCl facilitated the loss of bacterial culturablity more rapidly at the refrigerator temperature. However, it seemed likely clear that changes of the culturability are not the determination of making a decision on the true bacterial viability. In order to determine if bacterial cells are not dead, but still alive, it is inevitable to evaluate the degree by which organisms are sincerely damaged. Accordingly, a part of fluorescent probes can reflect the level of cellular integrities quantitatively. Recently, staining dyes of SYTO9 and propidium iodide (PI) are widely applied in a field of the microbiology because these dyes are well-known for the ability to penetrate inside the cellular membrane with increasing cell injuries. SYTO9 is capable of penetrating bacteria with the intact membrane, interacting with the cell nucleic acid, and then displaying green colors for the live cell with the fluorescence microscopy. Propidium iodide can only penetrate bacterial cells through damaged membranes, labeling the dead cell only as red-coloured fluorescence. Thus, staining bacterial cells with SYTO9 and propidium iodide can distinguish between live and dead cells. Loss of the permeability barrier of stained cells represents irreparable membrane damages and thus cell death (16-17). In the present study, three strains of *V. parahaemolyticus* showed the viable numbers in the levels of >5.5 log_10_ CFU per a slide uniformly until these pathogens were challenged by cold-starvation stress (nutrient-deprivation and cold temperature) for more than 100 days, regardless of the controlled environmental effector such as excessive concentrations of NaCl and acidic pHs, which means that although the culturable count of *V. parahaemolyticus* were below the detectable levels when incubated in ASW microcosms at 4°C for 80 days bacterial cells were proven to be still alive, maintaining the intact cell membrane. Accordingly, it was demonstrated that a prolonged period of cold-starvation challenge enabled three strains of *V. parahaemolyticus* to enter into the VBNC state within 80 days.

**FIG 1.**
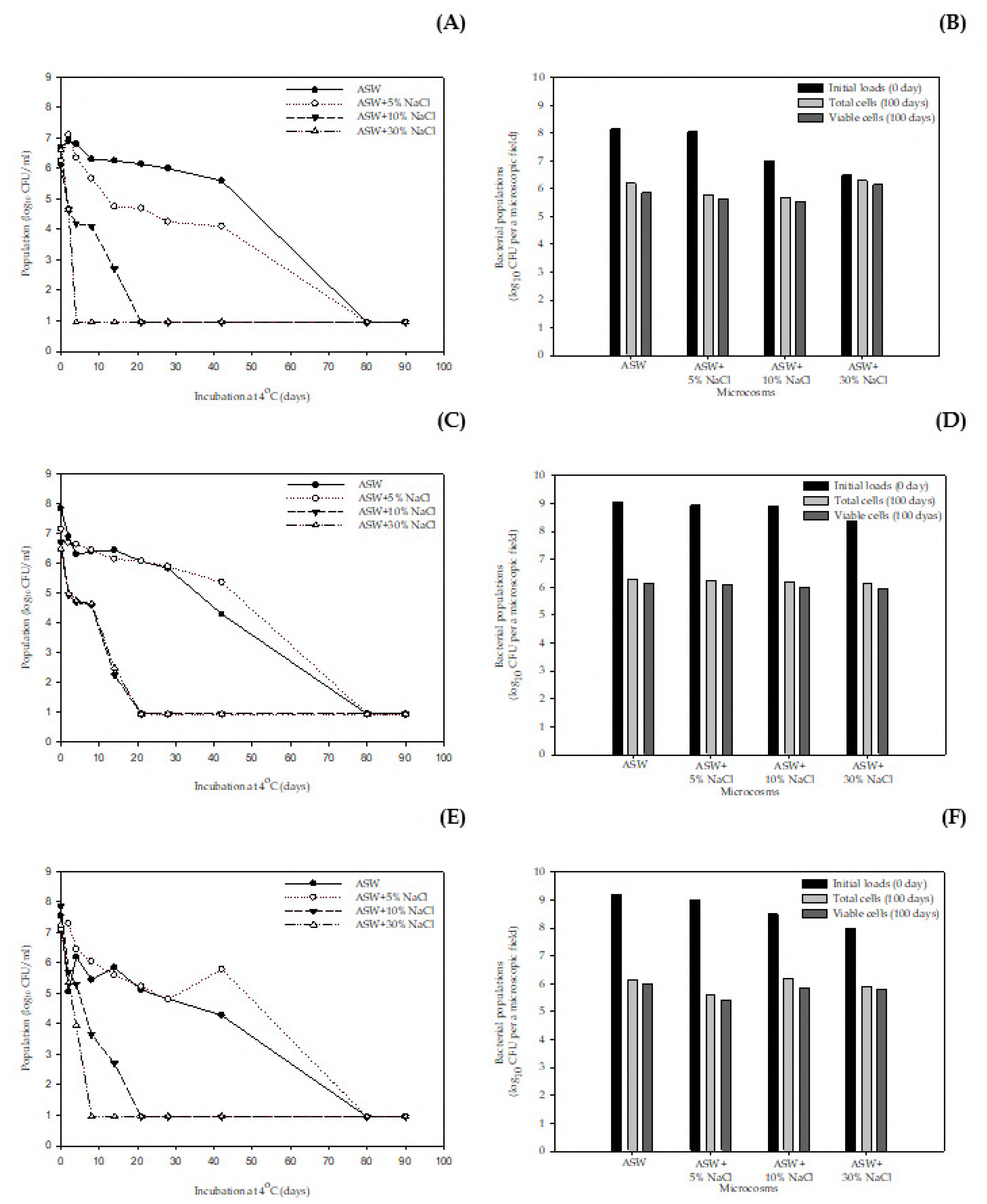
Loss of the culturability (A, C, and E) and the viability (B, D, and F) of *V. parahaemolyticus* ATCC 17082 (A-B), *V. parahaemolyticus* ATCC 33844 (C-D), and *V. parahaemolyticus* ATCC 27969 (E-F) incubated in ASW (pH 6) microcosms supplemented with varying concentrations of NaCl at 4°C for 100 days.

### Cytotoxic effects of VBNC pathogens

As shown in Fig. 2A, there was no difference in the cellular virulence (cytotoxicity) between the cells of *V. parahaemolyticus* ATCC 17082 either actively grown overnight in TSB at 37°C or incubated in ASW microcosms at 4°C for ≤80 days. In contrast, VBNC *V. parahaemolyticus* ATCC 17082 exhibited decreased cellular damages in the levels of ≤45% against Vero cells when incubated in ASW microcosms added with 5% NaCl at 4°C for 80 days previously (Fig. 2B). Cells of *V. parahaemolyticus* ATCC 33844 and *V. parahaemolyticus* ATCC 27969 showed gradual decreases in the cytotoxicity during cold-starvation. In particular, less than 40% of the cellular toxicity was achieved when these bacteria were incubated in ASW microcosms at 4°C for 21 days. Clearly, it appeared that the cellular virulence of *V. parahaemolyticus* ATCC 33844 and *V. parahaemolyticus* ATCC 27969 against Vero cells declined in the reverse proportion to a prolonged duration of cold-starvation. Especially, VBNC *V. parahaemolyticus* ATCC 33844 showed no cytotoxic effects on Vero cells as maintained in ASW microcosms supplemented with 30% NaCl at 4°C for 80 days (Fig. 2B). However, it was shown that once challenged by the cold-starvation for 80 days VBNC *V. parahaemolyticus* ATCC 27969 represented the strong cytotoxicity, disrupting the cellular structure of animal cell lines apparently (*data not shown*). Of much importance, strains of *V. parahaemolyticus* which were converted from the nonculturable state recovered the whole cytotoxicity in the levels of ≥100%. It has been well-reported that VBNC *V. parahaemolyticus* can be recovered back to the culturable state by eliminating the causative environmental stress. Several studies showed that strains of *V. parahaemolyticus* and *V. vulnificus* in such a dormant state were converted to the culturable state on solid agar plates, followed by culturing these long-term-stressed cells in liquid nutrient-rich media at ambient temperatures for several days (6, 35). Dwidjosiswojo et al. (18) reported that while VBNC-induced cells of *Pseudomonas aeruginosa* did not show the cytotoxicity these cells re-cultured in a nutrient-rich culture broth (which recovered back to the culturable state) killed 100% Chinese hamster ovary cells within 24 hrs, supporting these findings. According to Wong et al. (2), cells of *V. parahaemolyticus* were induced into the VBNC state when incubated in Morita minimal salt broth at 4°C for 35 days. Consequently, either VBNC or resuscitated cells of *V. parahaemolyticus* maintained higher levels of the cellular virulence, causing 100% cell death of Hep-2 cells within 24 hrs. Findings of Cappelier et al. (12) were also in an agreement with these results. Thus, it should be noted that VBNC cells of *V. parahaemolyticus* were not dead, but rather maintained peculiar metabolic activities to survive in an unfavorable environment such as cold-starvation. In this study, it was revelated out that three strains of *V. parahaemolyticus* in the VBNC state still remained the cellular toxicity against human cell lines, including Vero and CACO-2. Notably, VBNC *V. parahaemolyticus* strains were able to be resuscitated after temperature upshift, and then resulted in lethal injuries to the human cell lines. Therefore, it could be postulated that VBNC cells of pathogenic bacteria, including *V. parahaemolyticus,* would retain the infectious virulence or re-gain the infectivity after resuscitation process. VBNC-induced pathogen on marine products may present a potential risk of causing public health hazards. Necessarily, it should be further required for determining the environmental stress which activates either the transition of the food-borne pathogens into the VBNC state or the resuscitation-availability of VBNC bacteria to the culturable and infectious state.

**FIG 2.**
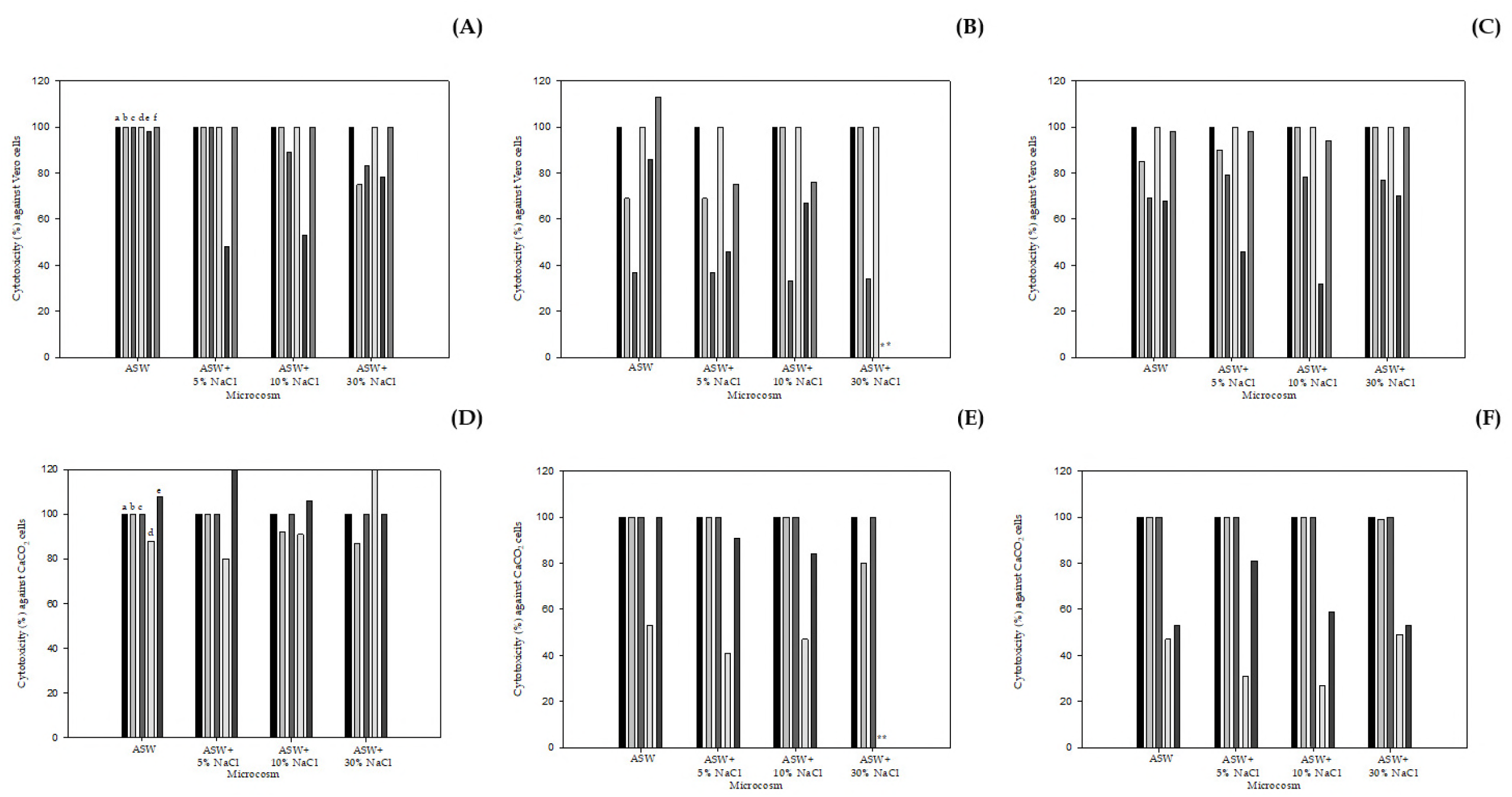
Changes of the cytotoxicity of *V. parahaemolyticus* ATCC (A and D) 17082, *V. parahaemolyticus* ATCC (B and E) 33844, and *V. parahaemolyticus* ATCC (C and F) 27969 incubated in ASW (pH 6) microcosms at 4°C for 80 days against (A-C) Vero and (D-F) CACO-2 cells, respectively. (A-C); a, 0 day; b, 7 days; c, 21 days; d, resuscitated cells in TSB after 21 days of cold-starvation; f, 80 days; g, resuscitated cells in TSB after 21 days of cold-starvation; *, not determined. (D-F); a, 0 day; b, 7 days, c, resuscitated cells in TSB after 21 days of cold-starvation; d, 80 days; e, resuscitated cells in TSB after 80 days of cold-starvation; *, not determined. Cells of VBNC *V. parahaemolyticus* suspended at 4°C for 21 and 80 days were transferred onto TSB, and then incubated at 25°C for consecutive 3 days, respectively.

### Assessment of cell membrane permeabilization

In the present study, inner and outer cell membrane permeabilization assays were carried out by using ß-galactosidase and ß-lactamase, respectively (Fig. 3). Outer cell membranes of *V. parahaemolyticus* were damaged under cold-starvation challenge at 4°C for 30 days as being assessed with the ß-lactamase activity. Commonly, optical densities at 482 nm of *V. parahaemolyticus* ATCC 17082 grown overnight in TSB and in the VBNC state were increased with the prolonged duration. In contrast, ß-galactosidase activities of VBNC *V. parahaemolyticus* were not different from the pure cultures of *V. parahaemolyticus* ATCC 17082. Given that ß-galactosidase is localized in cytoplasm and is capable of hydrolyzing ONPG if the inner membrane of bacteria is compromised (19), it seemed likely that such a cold-starvation stress interacted with the outer cell membrane either mainly or preferentially. In addition, ß-lactamase, which is present in the periplasmic space, responds to pyridine-2-azo-4’-(N’, N’-dimethylaniline) cephalosporin (PADAC) unless permeability of the outer membrane is decreasing (20). From these results, it could be accepted that the release of intracellular materials contributed to the loss of cell membrane permeabilization during a long-term of cold-starvation stress.

**FIG 3.**
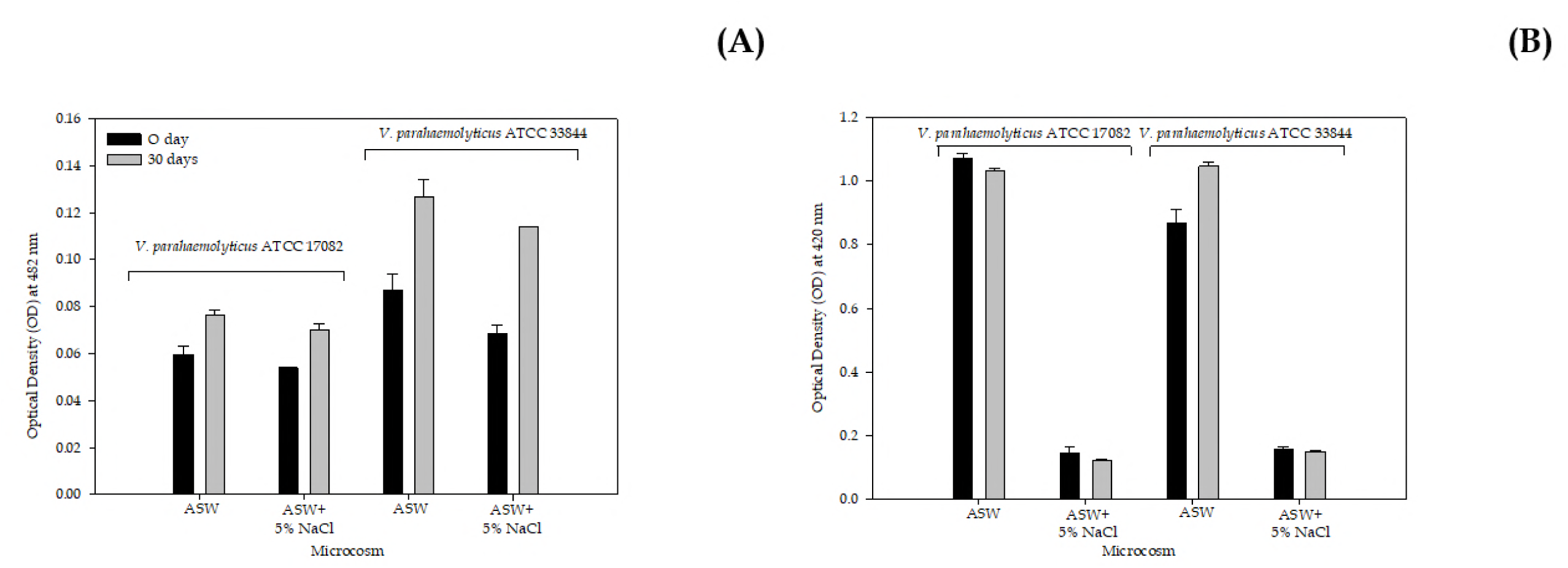
Assessment of the cellular membrane permeability (A, ß-lactamase activity; B, ß-galactosidase activity) of *Vibrio parahaemolyticus* ATCC 17082 and *V. parahaemolyticus* ATCC 33844 incubated in ASW microcosms at 4°C for 30 days.

The cellular leakage of nucleic acid and protein into microcosm fluids was also measured to evaluate the degree to which cell membrane permeabilization of VBNC *V. parahaemolyticus* was truly damaged. In Table 1, initial densities of nucleic acid ranged from 0.417 to 1.567 when *V. parahaemolyticus* began to be exposed at 4°C in ASW microcosms. After 150-days-incubuation at 4°C, higher levels of nucleic acid were released from these cells; leakages of nucleic acid were 2.611, 2.550, 2.315, and 2.683 from VBNC *V. parahaemolyticus* ATCC 33844 in ASW microcosms at controlled NaCl concentrations of 0.75%, 5%, 10%, and 30%, respectively. Similarly, it has been shown that intracellular proteins were permeabilized through the membrane of VBNC cells into bacterial microcosms. After the entry of *V. parahaemolyticus* into the VBNC state, nucleic acid and protein outflowed more than twice over those of the actively growing cells. These results implied that cold-starvation stress could increase the membrane permeability of *V. parahaemolyticus*, resulting in the cellular leakages of nucleic acid and protein from the bacterial cytoplasm to external fluids. According to Zhang et al. (21), these authors stated that the release of nucleic acid and protein would be strongly involved in persisting the integrity of cell membrane. At this point of view, the temporary accumulation of these cellular components in ASW microcosms was implicated with the loss of membrane integrity (22).

**TABLE 1.** Measurement of the cellular leakage (nucleic acid, at OD 260 nm) of *V. parahaemolyticus* cells incubated in ASW (pH 6) microcosms at 4°C for 150 days

| Pathogen | ATCC | Before (0 hrs): |  |  |  | After (150 days): |  |  |  |
| --- | --- | --- | --- | --- | --- | --- | --- | --- | --- |
|  |  | ASW | ASW+5% NaCl | ASW+10% NaCl | ASW+30% NaCl | ASW | ASW+5% NaCl | ASW+10% NaCl | ASW+30% NaCl |
| *V. parahaemolyticus* | 17082 | 0.433 | 0.502 | 0.598 | 0.974 | 0.868 | 1.200 | 0.999 | 1.053 |
| *V. parahaemolyticus* | 33844 | 0.576 | 0.685 | 0.783 | 1.562 | 2.611 | 2.550 | 2.315 | 2.683 |
| *V. parahaemolyticus* | 27969 | 0.751 | 0.778 | 0.659 | 1.107 | 3.255 | 3.417 | 3.385 | 3.755 |

### Enzymatic activities of catalase and GST

While culturing aerobically, reactive oxygen species (ROS) such as 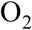, ^·^OH, and H_2_O_2_ are generally accumulated in the cell of bacteria. Especially, hydrogen peroxide (H_2_O_2_) has been recognized as one of the leading toxic compounds due to its ability to penetrate the cellular wall and membrane, thereby damaging nucleic acids directly (23). During exposure to environmental stressors such as starvation, bacteria might encounter intracellularly accumulated H_2_O_2_ (24). Hydrogen peroxidase is able to separate ROS into a molecule of H_2_O and O_2_. Generally, detoxifying enzymes, including catalase, GST, and peroxidase, prevented microbial cells from being damaged by ROS. It was determined that when *Ralstonia solanacearum* was induced into the VBNC state upon exposure to 200 ưM copper, significantly (p < 0.05) higher amounts of H_2_O_2_ were accumulated in the bacterial pellets (25). Such a cellular damage caused by H_2_O_2_ could have a strong influence on the entry of pathogens into the VBNC state. In particular, it has been reported that cells of *V. vulnificus* incubated in ASW microcosms added with ≤500 ưM H_2_O_2_ at 4°C were converted to the nonculturable state more rapidly (26). Thus, effects of H_2_O_2_ to the cellular damage could be examined by measuring the amounts of catalase produced intracellularly. In Fig. 4, *V. parahaemolyticus* ATCC 17082 grown in TSB exerted <20 U/mg/protein catalase, whereas VBNC-induced cells in ASW microcosms containing 0.75% NaCl and supplemented with 5%, 10%, and 30% NaCl showed <60, ≤100, <30, and <30 U/mg/protein catalase, respectively. When incubated in ASW microcosms amended with ≥10% NaCl at 4°C for 100 days *V. parahaemolyticus* ATCC 27969 exhibited a less decreased catalase activity (<50 U/mg/protein), as compared with that of the stationary-phase cells. While initial catalase activities of *V. parahaemolyticus* ATCC 33844 were <30 U/mg/protein, these organisms suspended in ASW microcosms added with ≥5% NaCl at 4°C for 100 days displayed enhanced catalase activities, ranging from 80 to <140 U/mg/protein. These results indicated that H_2_O_2_-degrading capability of *V. parahaemolyticus* would be dependent on strain-to-strain variables. GST was also recognized as one of the important agents associated with oxidative stress response of pathogens. Three strains of *V. parahaemolyticus* showed no differences among GST activities before and after a prolonged duration of cold-starvation (Fig. 5). Similarly, GST activities of these organisms ranged from >1.5 to <2.0 U/mg/protein. In the present study, there was no close correlation between the loss of culturability and ROS-detoxifying activities.

### Assessment of the Cellular Hydrophobicity

In Table 3, hydrophobicity of *V. parahaemolyticus* in the VBNC state at 4°C for 100 days was measured for determining a hallmark of VBNC-induced bacteria. When subjected to the refrigerator temperature for 100 days, levels of hydrophobicity of *V. parahaemolyticus* ATCC 17082 in ASW microcosms added with 5%, 10%, and 30% NaCl were 37.5%, 46.8%, and 62.2%, respectively. Surface hydrophobicity of VBNC *V. parahaemolyticus* ATCC 33844 and *V. parahaemolyticus* ATCC 27969 increased dramatically more than those of the pure cultures. Strains of *V. parahaemolyticus* used in this study exhibited the highest hydrophobicity of ≤62.2%, followed by cold-starvation in ASW microcosms supplemented with 30% NaCl at 4°C for 100 days. Probably, hydrophobicity of bacterial cells was closely related with a change of fatty acid composition, thereafter implying that the increasing amounts of saturated fatty acid (SFA) would result in a rise of the hydrophobicity of bacterial cells. Wong et al. (2) reported that total amounts of saturated fatty acid were largely declined in cells of *V. parahaemolyticus* ST550 during cold-starvation for 40 days, ranging from 60% to <40%. In this point of view, our results shown in Table 4 would indicate that decreases of the content of SFA, followed by a decline of the hydrophobicity, in VBNC cells of *V. parahaemolyticus* would correspond to the increasing resistances to environmental challenges. Several studies reported that nonculturable cells of bacteria, including *V. parahaemolyticus*, *V. vulnificus*, and *Campylobacter jejuni*, exerted notably higher resistances to 20 ppm H_2_O_2_, antibiotics, and heat (55°C for 3 min), supporting our findings (14, 27-28).

**TABLE 2.** Measurement of the cellular leakage (protein, at OD 280 nm) of *V. parahaemolyticus* cells incubated in ASW (pH 6) microcosms at 4°C for 150 days

| Pathogen | ATCC | Before (0 hrs): |  |  |  | After (150 days): |  |  |  |
| --- | --- | --- | --- | --- | --- | --- | --- | --- | --- |
|  |  | ASW | ASW+5% NaCl | ASW+10% NaCl | ASW+30% NaCl | ASW | ASW+5% NaCl | ASW+10% NaCl | ASW+30% NaCl |
| *V. parahaemolyticus* | 17082 | 0.390 | 0.435 | 0.465 | 0.797 | 0.426 | 0.602 | 0.600 | 0.638 |
| *V. parahaemolyticus* | 33844 | 0.466 | 0.537 | 0.630 | 1.316 | 1.270 | 1.218 | 1.226 | 1.433 |
| *V. parahaemolyticus* | 27969 | 0.573 | 0.597 | 0.521 | 0.910 | 1.588 | 1.747 | 1.872 | 2.351 |

**TABLE 3.**
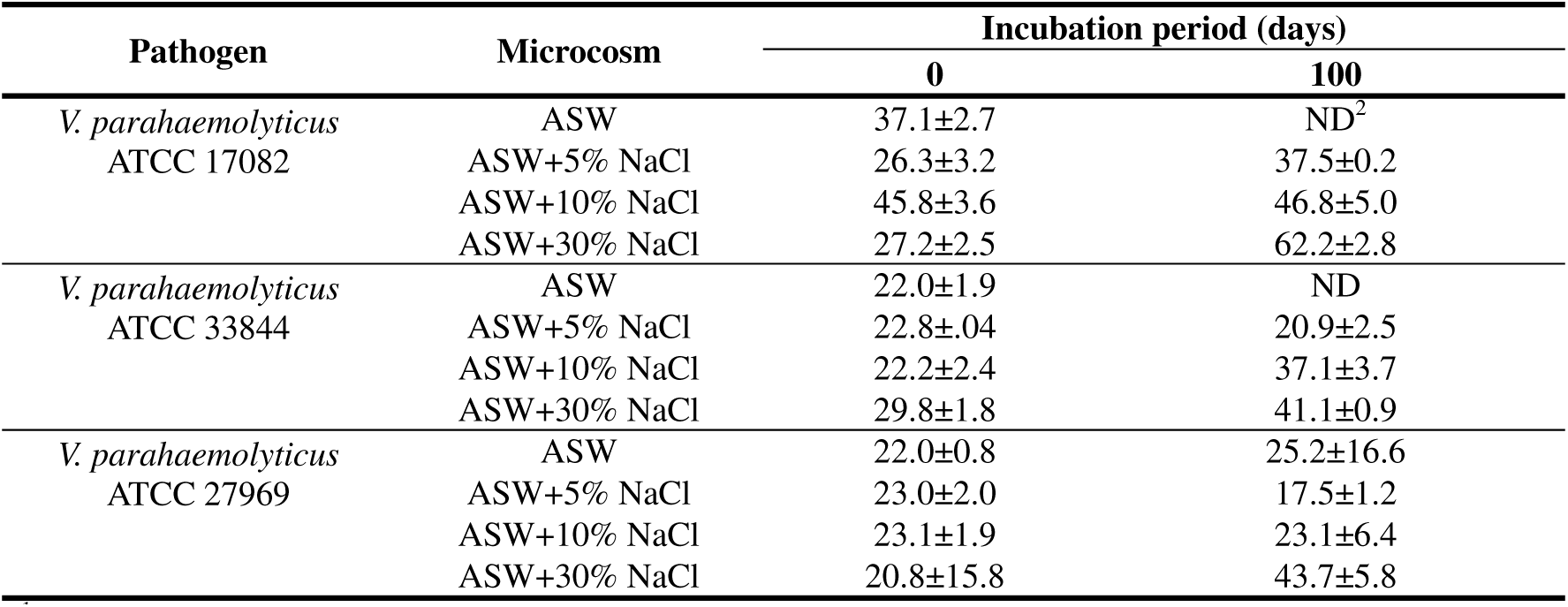
Measurement of hydrophobicity (%) of viable-but-nonculturable *V. parahaemolyticus* at 4°C for 100 days

**TABLE 4.** Comparison on the fatty acid composition (%) of viable-but-nonculturable *V. parahaemolyticus* ATCC 17082 at 4°C for 90 days

| Fatty acid | Name | Category | Microcosm |  |  |  |  |
| --- | --- | --- | --- | --- | --- | --- | --- |
|  |  |  | Ctrl | ASW | ASW+5% NaCl | ASW+10% NaCl | ASW+30% NaCl |
| 11:0 iso 3OH | 3-Hydroxy-9-Methyldecanoic acid | Branched-chain | 0.75% | ND<sup>1</sup> | ND | 0.30% | 0.32% |
| 12:0 | Lauric acid | Saturated | 2.02% | 4.37% | 4.38% | 4.37% | 4.44% |
| 12:0 2OH | 2-Hydroxylauric acid | Hydroxy | 1.63% | 1.94% | 3.12% | 3.05% | 2.92% |
| 14:0 | Myristic acid | Saturated | 7.25% | 8.05% | 9.11% | 9.19% | 8.81% |
| 14:0 3OH | ND | Hydroxy | 1.86% | 1.50% | 1.68% | 1.58% | 1.84% |
| 16:0 | Palmitic acid | Saturated | 25.64% | 28.65% | 23.88% | 23.36% | 23.05% |
| 16:0 N alcohol | Cetyl alcohol | Saturated | -<sup>2</sup> | ND | 0.20% | 0.40% | 0.50% |
| 18:0 | Stearic acid | Saturated | 2.19% | 2.71% | ND | 1.23% | 1.22% |
| Total (saturated fatty acid) |  |  | 41.34% | 47.22% | 42.37% | 43.48% | 43.10% |
| 16:1 w7c | Palmitoleic acid | Unsaturated | 26.76% | 24.52% | 32.98% | 34.11% | 35.77% |
| 16:1 w6c | cis-10-Palmitoleic acid | Unsaturated | 3.78% | 7.13% | 5.11% | 4.44% | 3.82% |
| 16:1 w5c | cis-11-Palmitoleic acid | Unsaturated | - | - | 0.50% | 0.42% | 0.53% |
| 16:1 w7c alcohol | Palmitoleyl alcohol | Unsaturated | - | - | - | 0.37% | - |
| 17:1 anteiso w9c | (7Z)-13-Methyl-7-Hexadecenoic acid | Unsaturated | 7.52% | 3.12% | 2.13% | 2.42% | 2.84% |
| 17:1 iso | Isomargaric acid | Branched-chain | - | - | 0.85% | 0.24% | - |
| 18:1 w9c | Oleic acid | Unsaturated | - | - | - | 0.38% | - |
| 18:1 w7c | cis-Vaccenic acid | Unsaturated | 20.61% | 18.01% | 14.90% | 14.16% | 13.18% |
| 18:1 w5c | cis-13-oleic acid | Unsaturated | - | - | - | 0.35% | - |
| 18:2 w6c | Linoleic acid | Unsaturated | - | - | - | - | 0.37% |
| Total (unsaturated fatty acid) |  |  | 58.67% | 52.78% | 56.47% | 56.89% | 56.51% |

<sup>1</sup> ND, not determined.<sup>2</sup> -, not found.

**FIG 4.**
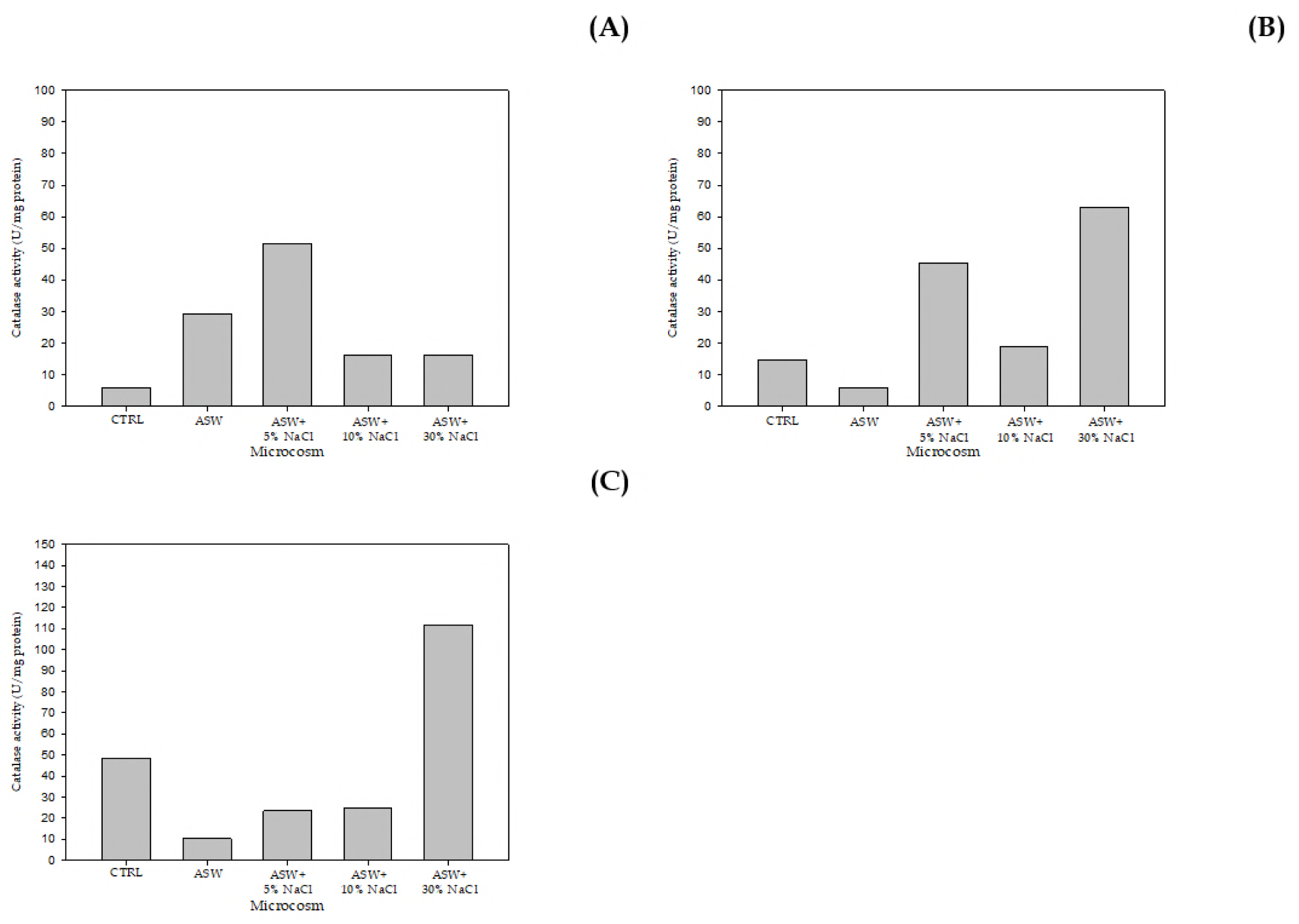
Assessment of catalase activity (U/mg protein) of *V. parahaemolyticus* ATCC (A) 17082, *V. parahaemolyticus* ATCC (B) 33844, and *V. parahaemolyticus* ATCC (C) 27969 in the VBNC state at 4°C for 100 days.

**FIG 5.**
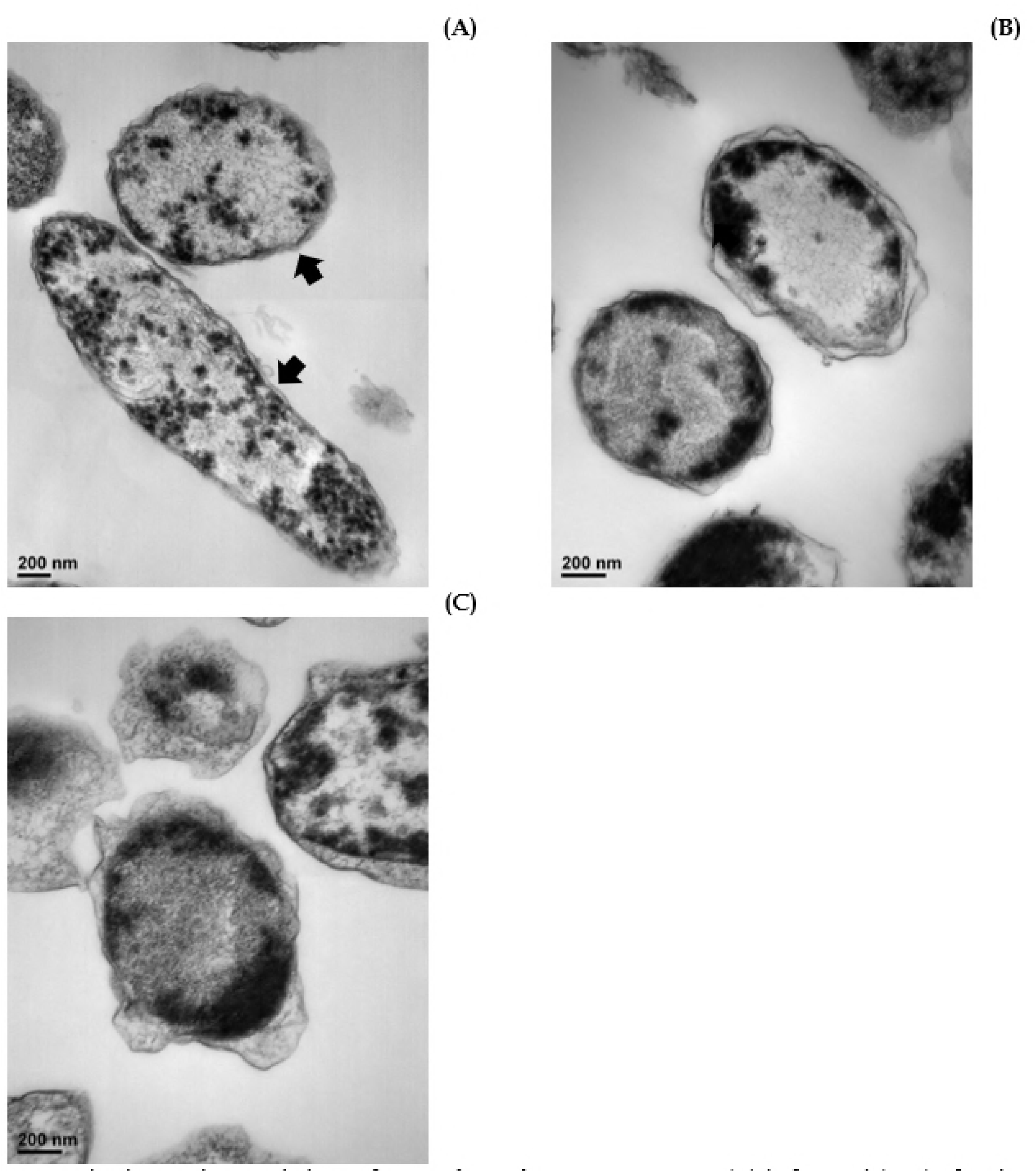
Investigation on the morphology of *V. parahaemolyticus* ATCC 33844 (A) before and (B-C) after the entry into the VBNC state. Bacterial cells of *V. parahaemolyticus* ATCC 33844 were incubated in ASW (pH 6.1) microcosms containing (B) 0.75% NaCl or (C) supplemented with 5% NaCl at 4°C for 100 days.

### Morphological changes of VBNC cells

In a stationary-phase grown cell of *V. parahaemolyticus* ATCC 17082 (Fig. 6A), the cytoplasmic matrix was filled with granules and the cell membrane became intact without minor damages. On contrast, VBNC cells of *V. parahaemolyticus* ATCC 17082 were shown to have less densed cytoplasmic spaces than those of the stationary-phase cells. In particular, cellular membranes of VBNC cells were largely loosed, resulting in generating empty spaces between the outer and inner membranes (Fig. 6B-C). When it comes to a study conducted by Zhao et al. (29), it was reported that the number of ribosomes was notably reduced in VBNC cells of *Escherichia coli* O157:H7. These authors also pointed out that this change in the interior structure of cells would be implicated with decreases of both deoxyribonucleic acid (DNA) amplification and protein translation, thereby causing these VBNC cells to minimize cell maintenance requirements (compactness)

**FIG 6.**
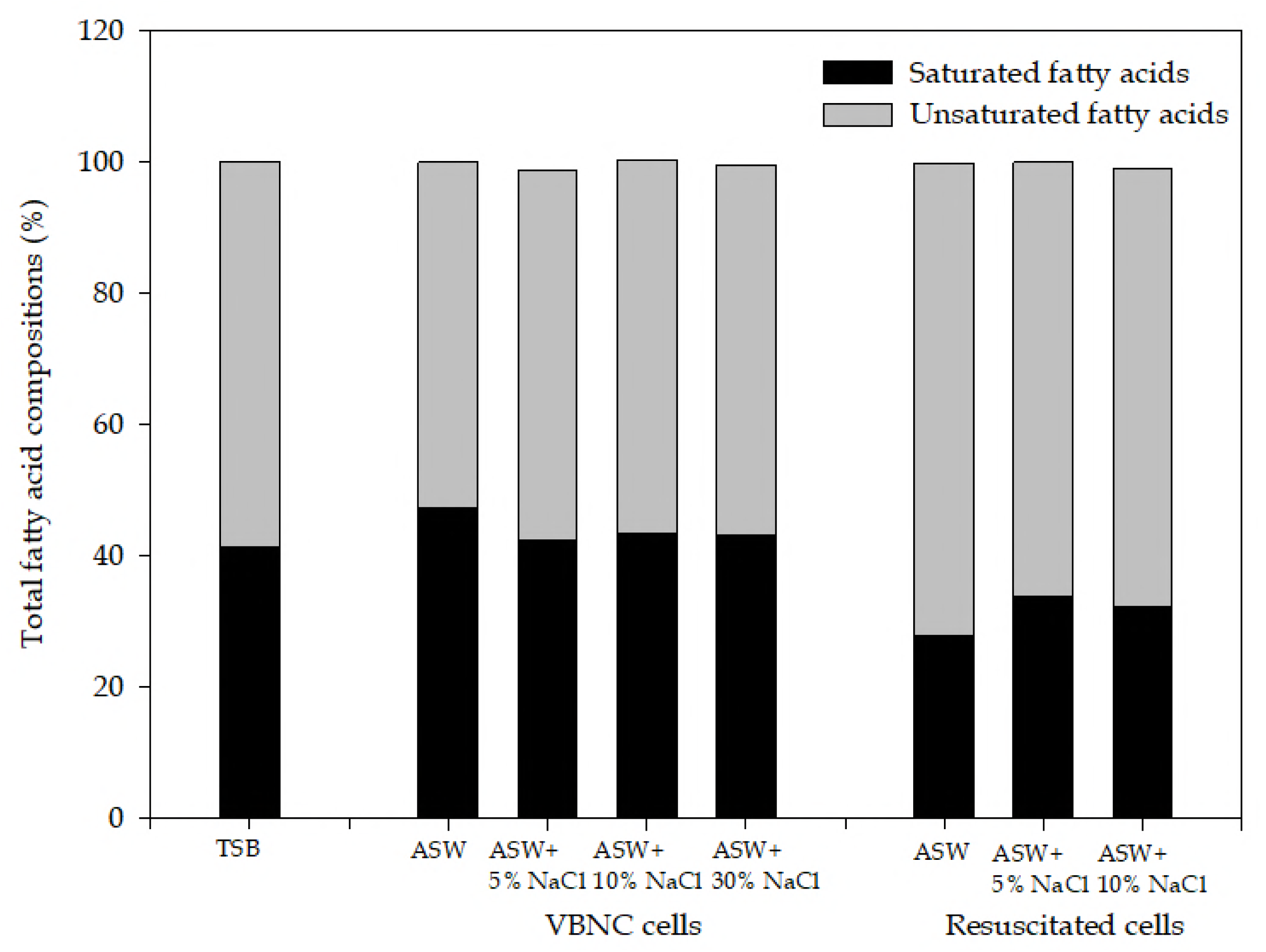
Comparison of fatty acid compositions between *V. parahaemolyticus* ATCC 17082 incubated in ASW (pH 6) microcosms at 4°C for 90 days and resuscitated cells.

### Fatty acid compositions of VBNC *V. parahaemolyticus*

Cells of *V. parahaemolyticus* ATCC 17082 grown overnight in TSB accounted for 41.34% of the saturated fatty acid and 58.67% of unsaturated fatty acid out of the whole fatty acid composition. Normal cells exerted the highest amounts of palmitic acid (C_16:0_, 25.64%) and palmitoleic acid (C_16:1 w7c_, 26.76%), respectively. When cells of *V. parahaemolyticus* ATCC 17082 were incubated in ASW microcosms added with higher concentrations of NaCl at 4°C for 90 days, contents of palmitic acid were increased slightly, ranged from 23.05% to 23.88%. Among the saturated fatty acid, contents of lauric acid (C_12:0_), 2-hydroxylauric acid (C_12:0 2OH_), and myristic acid (C_14:0_) were increased as *V. parahaemolyticus* ATCC 17082 became the viable-but-nonculturable state. In terms of the unsaturated fatty acid, VBNC cells exhibited increasing concentrations of palmitoleic acid, ranging from 32.98% to 35.77%. In particular, it was shown that unsaturated fatty acids were more synthesized with the increasing concentrations of NaCl. On the other hand, levels of cis-vaccenic acid (C_18:1 w7c_) declined largely upon exposure of *V. parahaemolyticus* to the low temperature for 90 days in ASW microcosms with the addition of increasing amounts of NaCl, showing by 18.1%, 14.90%, 14.16%, and 13.18%, respectively. In addition, it seemed clear that several saturated fatty acids, including lauric acid, 2-hydroxylauric acid, and myristic acid, were responsible for the increasing amounts of total saturated fatty acid in VBNC cells of *V. parahaemolyticus* ATCC 17082. In a study conducted by Jia et al., (30), four strains of *V. parahaemolyticus* isolated from seafood showed that palmitic acid (C_16:0_) was decreased dramatically, ranging from 47.6% to 9.9% when these organisms were incubated in TSB at 4°C for 30 days, whereas palmitoeleate (C_16:1_) increased greatly after the entry into the VBNC state. This study was in an agreement with our findings. It was also reported that palmitic acid contents (%) of *V. vulnificus* C7184 were significantly (p < 0.05) lower once this organism was incubated in ASW microcosms at 5°C for 24 hrs (31). In VBNC cells of *V. vulnificus*, palmitoleic acid (C_16:1_) and stearic acid (C_18:0_) increased significantly (p < 0.05), while margaric acid (C_17:0_) declined remarkably at the refrigerator temperature within 24 hrs, supporting our findings. Furthermore, it was demonstrated that VBNC cells of *V. parahaemolyticus* ATCC 17082 were recovered to the culturable state reversibly as being resuscitated in a nutrient-rich medium such as TSB (pH 8) and brain hearth infusion broth (pH 8) at 25°C for several days (*data not shown*). Thus, resuscitated cells were further used to perform the fatty acid composition analysis in the present study. As shown in Table 6, the total density of unsaturated fatty acids was increased more than normal and VBNC cells. Especially, lauric acid (C_12:0_), 2-Hydroxylauric acid (C_12:0 2OH_), and unknown (C_14:0 3OH_) were increased comparatively, whereas palmitic acid (C_12:0_) was decreased remarkably among the total saturated fatty acid of resuscitated cells. After resuscitation, cis-vaccenic acid (C_18:1 w7c_) increased largely, ranging from 20.65% to 32.17%. Of which on the resuscitated cells of *V. parahaemolyticus* ATCC 17082, the total amounts of saturated fatty acid declined, but unsaturated fatty acids increased to that extent. In our preliminary studies, VBNC *V. parahaemolyticus* exhibited higher resistances through a simulated human gastric digestion (*data not shown*). Along with our findings in this study, it would be suggested that increasing synthesis of saturated fatty acids in the VBNC cells correspond to the improved environmental resistance. As saturated fatty acids display a straight-line structure VBNC cells would acquire more compact cell membranes, thereby resulting in more decreased cellular permeabilization than that of the normal cells. Consequently, it would administer higher resistances to various environmental challenges for these VBNC cells. Previously, several studies disclosed that there were significantly different fatty acid compositions on the bacterial membranes between actively growing and VBNC cells (2, 31). Interestingly, it has been shown that ratios of saturated fatty acids/unsaturated fatty acids increased significantly after forming VBNC cells. Probably, this change would be closely associated with a distinct physiology of VBNC cells. Given that VBNC cells might be capable of re-gaining the platable-capability on culture media, followed by further resuscitation efforts, there are still restricted information available on understanding cellular properties of VBNC *V. parahaemolyticus*. By comparing the fatty acid composition between VBNC and resuscitated cells, it would provide a meaningful insight in revealing out mechanisms governing the entry of *V. parahaemolyticus* into the VBNC state.

**TABLE 5.** Comparison of fatty acid composition (%) of *V. parahaemolyticus* ATCC 17082 to be recovered from the VBNC state

| Fatty acid | Name | Category | Microcosm |  |  |  |
| --- | --- | --- | --- | --- | --- | --- |
|  |  |  | Ctrl | ASW | ASW+5% NaCl | ASW+10% NaCl |
| 11:0 iso 3OH | 3-Hydroxy-9-Methyldecanoic acid | Branched-chain | 0.75% | 0.39% | -<sup>1</sup> | - |
| 12:0 | Lauric acid | Saturated | 2.02% | 2.71% | 2.75% | 2.95% |
| 12:0 2OH | 2-Hydroxylauric acid | Hydroxy | 1.63% | 2.48% | 2.55% | 2.78% |
| 13:0 iso | Isotridecanoic acid | Branched-chain | - | 0.65% | 0.63% | 0.65% |
| 13:0 3OH | ND | Branched-chain | - | - | 0.29% | 0.28% |
| 14:0 | Myristic acid | Saturated | 7.25% | 4.50% | 4.18% | 4.70% |
| 14:0 3OH | ND | Hydroxy | 1.86% | 2.79% | 3.09% | 2.40% |
| 15:0 anteiso | 12-Methylmyristic acid | Saturated | - | - | - | 0.38% |
| 16:0 | Palmitic acid | Saturated | 25.64% | 10.40% | 15.52% | 14.65 |
| 16:0 iso | Isopalmitic acid | Branched-chain | - | - | 0.33% | 0.52% |
| 16:0 N alcohol | Cetyl alcohol | Saturated | - | - | - | - |
| 17:0 | Margaric acid | Saturated | - | 1.45% | 2.11% | 1.75% |
| 17:0 anteiso | Anteisomargaric acid | Branched-chain | - | 0.82% | 0.47% | 0.63% |
| 17:0 iso | Isomargaric aid | Branched-chain | - | - | 0.34% | 0.44% |
| 18:0 | Stearic acid | Saturated | 2.19% | 1.68% | 1.57% | 0.55% |
| Total (saturated fatty acid) |  |  | 41.34% | 27.87% | 33.83% | 32.30% |
| 16:1 w7c | Palmitoleic acid | Unsaturated | 26.76% | 24.55% | 23.14% | 26.80% |
| 16:1 w7c alcohol | Palmitoleyl alcohol | Unsaturated | - | - | 0.22% | - |
| 16:1 w6c | cis-10-Palmitoleic acid | Unsaturated | 3.78% | 7.03% | 8.22% | 10.16% |
| 17:1 anteiso w9c | (7Z)-13-Methyl-7-Hexadecenoic acid | Unsaturated | 7.52% | 3.83% | 1.09% | 2.38% |
| 17:1 w8c | cis-margaroleic acid | Unsaturated | - | 2.00% | 1.90% | 1.84% |
| 17:1 w6c | (11Z)-11-Heptadecenoic acid | Unsaturated | - | 1.39% | 1.22% | 1.37% |
| 18:1 w7c | cis-Vaccenic acid | Unsaturated | 20.61% | 32.17% | 30.38% | 23.98% |
| 18:2 w6c | Linoleic acid | Unsaturated | - | 0.93% | - | 0.26% |
| Total (unsaturated fatty acid) |  |  | 58.67% | 71.90% | 66.17% | 66.79% |

<sup>1</sup> -, not found.

**TABLE 6.** Determination of cellular properties of *V. parahaemolyticus* in the VBNC state

| Property<sup>1</sup> |  | Category |  |  |
| --- | --- | --- | --- | --- |
|  |  | Normal cells | VBNC cells | Resuscitated cells |
| Cell size |  | - | ▼ | - |
| Morphology |  | Rod | Coccus | Rod |
| Membrane damage |  | - | Blebbing | - |
| Cytotoxicity |  | ▲ | ▼ | ▲ |
| Hydrophobicity |  | - | ▲ | ND<sup>2</sup> |
| Fatty acid composition | SFA | - | ▲ | ▼ |
|  | USFA | - | ▼ | ▲ |
| Enzyme activity | Catalase | - | - | ND |
|  | GST | - | - | - |
| Cellular leakages |  | - | ▲ | ND |

<sup>1</sup> SFA, saturated fatty acid; USFA, unsaturated fatty acid; GST, glutathione-S-transferase.<sup>2</sup> ND, not determined.

Based on all the findings obtained from the present study, a novel hypothesis explaining the entrance of *V. parahaemolyticus* into the viable-but-nonculturable state would be established (Fig. 7). It would be plausible that the morphological transformation of bacterial cells either from rods to coccus or the dwarfing in the cellular size has to be preceded before the actively growing cells could turn into the VBNC state. Previously, it was determined that 100-days-stressed cells of VBNC. *V. parahaemolyticus* displayed the increasing cellular leakages such as nucleic acid and protein as much twice as the pure cultures. Besides, it was also shown that VBNC *V. parahaemolyticus* exhibited the increasing permeabilization through the outer membrane, indicating that the loss of cell membrane integrity caused by the increasing membrane permeabilization could result in the cellular leakage before VBNC cells could be steadily developed. Hence, such a transient conversion of bacterial cells could aim to alter the surface area as compact as possible, minimizing the cellular damage and the consumption of energy sources under continuous environmental stresses (nutrient-deprivation and cold temperature). One of our findings about the morphological change of VBNC *V. parahaemolyticus* can be in an agreement with this hypothesis strongly. Subsequently, fatty acid composition of these cells will begin to be altered, showing the increasing levels of saturated fatty acid. This procedure might be significantly associated with the complete compactness of the cellular membrane structure, and then could diminish the increased permeabilization of outer membranes into normal ranges, thereby stabilizing VBNC cells at the end. Seriously, as most VBNC cells of bacteria exerted higher resistances to various environmental conditions more than the actively grown cells enhanced survivals of VBNC cells could be elucidated by the alternation of fatty acid compositions after forming VBNC cells. Overall, it can be defined that once challenged by environmental stresses nonculturable bacteria, which show cellular modifications such as the increasing synthesis of saturated fatty acid, causing the cellular dwarfing in sizes to minimize the exposure of bacterial surfaces and to save the energy consumption, and then prevent intracellular periplasms from being seriously damaged by ROS, maintaining the integrity of bacterial membranes, could be called as viable-but-nonculturable cells.

**FIG 7.**
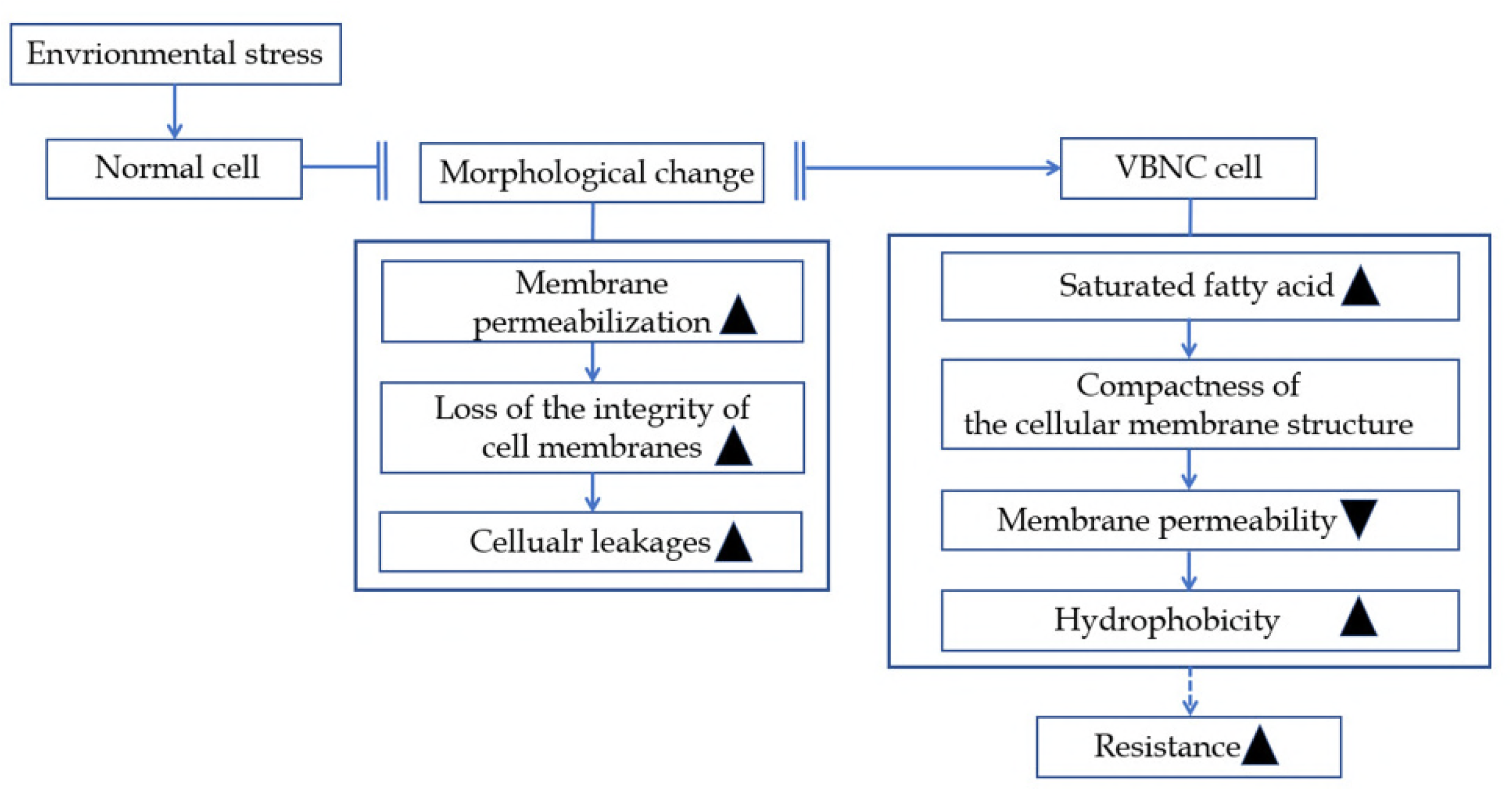
A schematic diagram for elucidating the transition process of *V. parahaemolyticus* cells into the viable-but-nonculturable state.

In conclusion, this research was conducted to determine the cellular characteristics of VBNC *V. parahaemolyticus*. As shown in Table 4, three strains of *V. parahaemolyticus* used in this study showed that VBNC cells (Ι) ratained the cellular virulence to Vero and CaCO_2_ cell lines, (ΙΙ) re-gained the 100% cytotoxicity even after resuscitation process, (ΙΙΙ) became permeabilized mainly at the binding sites of the outer membrane, (ΙV) showed lower levels of enzymatic (catalase and GST) activities, (V) exerted the increasing hydrophobicity, (VΙ) showed increasing amounts of SFA, and then (VΙΙ) displayed flappy outer membranes out of the cytoplasm in the cellular morphology. As pathogens can encounter a number of environmental stresses through a wide variety of channels contaminated food materials are subjected to heat, desiccation, cold temperature, freeze, and others in the middle of the preparation and the processing. As one of the surviving strategies, pathogens on food respond to these challenges, could turn into the VBNC state, and then can pose a risk of causing the food-borne outbreaks potentially. Clearly, it was demonstrated that VBNC cells could be the source of food-borne outbreaks upon eliminating the initiative environmental stress. Therefore, understanding the cellular characterization of nonculturable pathogens is of much importance in an attempt to designing adequate control measures for assuring the food safety. Although these results cannot reveal out all of the properties of VBNC bacteria, it will provide useful basic natures of a bacterial surviving strategy, understanding a mechanism governing the transition into the VBNC state.

## MATERIALS AND METHODS

### Preparation of microcosm solutions

ASW solutions (Sigma-Aldrich, St. Louis, MO, USA) were prepared according to the manufacturer’s instruction. Formal ASW microcosms (pH 7.2-7.8) contained 19,290 mg of Cl, 10,780 mg of Na, 2,660 mg of SO_4_, 420 mg of K, 400 mg of Ca, 200 mg of CO_3_, 8.8 mg of Sr, 5.6 mg of B, 56 mg of Br, 0.24 mg of I, 0.3 mg of Li, 1.0 mg of F, and 1,320 mg of Mg per 1 liter of sterile distilled water. These microcosms were autoclaved at 125°C for 20 min before use. Then, NaCl concentrations of these microcosms were adjusted to 0.75%, 5%, 10%, and 30% (m/v), respectively. Each of these microcosms was adjusted to pH 6.0–6.2, using a membrane-filtered 1N NaOH solution (Kanto chemical, Tokyo, Japan), facilitating the induction of *V. parahaemolyticus* into the VBNC state.

### Preparation of bacterial inoculums

Cells of *V. parahaemolyticus* ATCC 17082, *V. parahaemolyticus* ATCC 27969, and *V. parahaemolyticus* ATCC 33844 were purchased from the Korean Collection for Type Cultures (KCTC, Daejon, Korea). Bacterial stocks were maintained at −.75°C and further activated by a series of transfers in tryptic soy broth (Difco, Detroit, MI, USA) supplemented with 3% NaCl (TSB) at 37°C for 24 hrs before use. Each strain of *V. parahaemolyticus* ATCC 17082, *V. parahaemolyticus* ATCC 33844, and *V. parahaemolyticus* ATCC 27969 grown overnight in 35 ml of TSB at 37°C for 24 hrs was harvested by the centrifugation at 10,000 × g for 3 min, washed twice, and then bacterial pellets were re-suspended in 1 ml of ASW solutions (pH 6), corresponding to the bacterial density of approximately 10^8-9^ CFU/ml. Finally, cells of these pathogens were inoculated in 100 ml of ASW microcosms (pH 6) added with 0.75%, 5%, 10% and 30% NaCl, respectively. Bacterial fluids were kept at 4°C and ASW microcosms were withdrawn from the incubator at regular time-intervals to conduct the following process.

### Enumeration of the bacterial population

In order to count the bacterial number, cells of *V. parahaemolyticus* ATCC 17082, *V. parahaemolyticus* ATCC 33844, and *V. parahaemolyticus* ATCC 27969 were plating-counted on typtic soy agar (Difco) supplemented with 3% NaCl (TSA). Decimal dilutions (10^−1^) of these cells were prepared in alkaline peptone water (APW, pH 8) consisting 10 g of peptone and 10 g of NaCl per 1 liter of distilled water and 100 µl of these aliquots was spread on TSA. These media were further incubated at 37°C for 24 hrs. Colonies developed on media were further enumerated. At the same time, the numbers of living and dead cells of *V. parahaemolyticus* were measured using the Live/Dead^(R)^ BacLight Bacterial Viability kit (Life technologies, Eugene, Oregon, USA). Briefly, equal volumes (1:1) of SYTO9 and propidium iodide (PI) were combined and 3 µl of this mixture was added to each 1 ml of the bacterial cell. After a short time of incubation (approximately 15 min) at an ambient temperature in the dark, 5-8 µl of this aliquot was attached on a glass slide and a coverslip was placed on it. Then, bacterial images were demonstrated using an electron microscope (TE 2000-U, Nikon, Japan).

### Preparation of human cell lines

CACO-2 and Vero cells were cultured in 5-10 ml of Dulbecco’s modified eagle’s medium (DMEM, Corning, NY, USA) supplemented with 5% and 20% fetal bovine serum (FBS, Corning) at 37°C for 1-2 days in 5% CO_2_, respectively. The medium was removed in a petri-dish and washed 5 ml of PBS three times, and then 5 ml of trypsin (Corning) was added to facilitate the cell lysis. Each of these cells was incubated at 37°C for 5 min in 5% CO_2_. To alleviate the enzymatic activity caused by trypsin, 2-3 ml of the medium was added to all the cell cultures. Cell fluids were further transferred into 15 ml of a sterile cap tube and centrifugated at 15,000 × g for 3 min. The supernatant was eliminated, re-suspended in 5 ml of the medium, corresponding to the cellular density of 10^4^/ml, and then 100 μl of the eukaryotic cell solution was loaded into a 96-well plate containing 100 μl of the DMEM medium. CACO-2 and Vero cells were incubated at 37°C for 24 hrs in 5% CO_2_ before use.

### Measurement of the cytotoxicity of VBNC *V. parahaemolyticus* against the cell line

Cellular virulence of *V. parahaemolyticus* in the VBNC state against the human cell line was measured according to several studies conducted by Beattie and Williams (32), Jeßberger et al. (33), and Dektas et al. (34). As mentioned above, prepared cell lines, corresponding to 10^4^ CFU/ml, were added into a 96-well plate, respectively. Serial dilutions (0.1 ml/well) of the filtered cell-free bacterial supernatants were placed into 96-well plates. Or bacterial suspensions, corresponding to 10^8^ CFU/ml, was added in each of wells. Animal cell suspensions (0.1 ml/well), corresponding to 10^4^ cells per a well for Vero cells and 2 × 10^4^ cells per a well for CACO-2 cells, were supplemented into 96-well plates, respectively. Cell fluids were incubated at 37°C for 24 hrs under the atmospheric environment containing 5% CO_2_, replaced with either 100 μl/well of EMEM containing 5 mg/ml 3-(4, 5-dimethylthiazol-2-yl)-2, 5 diphenyl tetrazolium bromide (MTT, Corning), and then incubated at 37°C for 1 hr.

These mixtures were further removed in each of the wells, 100 μl of dimethyl-sulphoxide (DMSO, Corning) was added into a 96-well plate, and then read on a microtiter plate reader at 570 and 620 nm (Multiskan GO Microplate Spectrophotometer, Thermo Scientific, Vantaa, Finland).

### Analysis of the cell membrane permeabilization

The release of cytoplasmic ß-galactosidase from cells of VBNC *V. parahaemolyticus* was measured to determine the inner membrane permeabilization. Briefly, 10 μl of VBNC *V. parahaemolyticus* cells in ASW microcosms (pH 6) were transferred onto each well of a 96-well plate containing 50 μl of *O*-Nitrophenyl-_L_-_D_-galactoside (Sigma) and 40 μl of ASW solutions (pH 6). Once bacterial suspension was incubated at 30°C for 30 min in a dark room the absorbance was read on a microtiter plate reader at 420 nm (Multiskan GO Microplate Spectrophotometer). When it comes to determine the outer membrane permeabilization of VBNC cells, 10 μl of VBNC *V. parahaemolyticus* cells in ASW microcosms were added to each well of a 96-well plate containing 80 μl of ASW solutions and 10 μl of nitrocefin (Sigma), incubated at 30°C for 30 min in a dark room, and then the absorbance was read on a microtiter plate reader at 482 nm (Thermo Scientific).

### Analysis of the cellular leakage in the bacterial suspension

Cell membrane integrity was assessed by measuring cellular leakages such as nucleic acid and protein from VBNC cells of V. *parahaemolyticus*. Cell suspensions (1.5 mL) of *V. parahaemolyticus* ATCC 17082, *V. parahaemolyticus* ATCC 33844, and *V. parahaemolyticus* ATCC 27969 which had been incubated in ASW microcosms at 4°C for 100 days were transferred onto a sterile microtube, respectively. Pure cultures of these pathogens grown overnight in TSB at 37°C were centrifugated at 13,000 × g for 3 min, washed twice, and then re-suspended in ASW solutions (pH 8) added with 0.75%, 5%, 10%, and 30% NaCl, respectively. These pathogens were centrifuged at 10,000 × g at 4°C for 15 min to collect bacterial pellets. Each of these supernatants was collected in a sterile microtube and further used to assess the cellular leakage such as nucleic acid and protein by measuring the absorbance at 260 and 280 nm via multi-scan Go spectrophotometer (Thermo Scientific Inc.).

### Measurement of enzymatic activities

Catalase activity was measured in terms of H_2_O_2_-degradation with a spectrophotometric assay (CAT100, sigma). This assay was undertaken following by a method introduced in several studies (Santander et al., 2017; Molina-Cruz et al., 2008). Overnight grown cultures of *V. parahaemolyticus* and VBNC cells incubated in ASW (pH 6) microcosms at 4°C for 100 days were re-suspended in 50 mM potassium phosphate buffer (pH 7) containing 1 g of 3-mm-sized glass bead (Sigma), vortexed for 25 min, and then centrifugated at 13,000 × g for 3 min. According to a manual provided by a manufacture, supernatant was separately transferred into a sterile microtube. In a total of 100 μl of volume, 15 μl of supernatant was combined with 5 mM H_2_O_2_. Such a reaction was carried out at 25°C for 15 min and ceased with 900 μl of 15 mM sodium azide. Then, absorbance was colorimetrically read at 520 nm via multi-scan Go spectrophotometer (Thermo Scientific Inc.). Glutathione-S-Transferase (GST) activity was measured, using GST assay kit (CS0410, Sigma). In this assay, 1-Chloro-2, 4-DiNitroBenzene (CDNB) was used as a substrate. Twenty μl of bacterial supernatants were added to 180 μl of a substrate solution in each well of a 96-well plate, respectively. Then, absorbance was read at 340 nm via multi-scan Go spectrophotometer.

### Microbial adhesion to hydrocarbons (MATHs) assay

Cells of *V. parahaemolyticus* grown overnight in TSB and incubated in ASW microcosms at 4°C for 50 days were centrifugated at 8,000 × g for 1 min, washed twice, and then re-suspended in 0.1 M PBS to fit an optical density of 1.0 (A_o_) at 600 nm via a UV-Visible Spectrometer (Multiskan GO, Thermo Scientific). One hundred µl of hexadecane was added to 1 ml of these bacterial suspensions and incubated at an ambient temperature for 10 min. Optical density of these bacteria in the aqueous phase was measured at 600 nm (A_1_). The degree of hydrophobicity was calculated, following as (Choi, et al., 2013);

[1-A_1_/A_o_] × 100 (%).

### Transmission electron microscopic (TEM) Assay

Bacterial cells incubated in ASW microcosms (pH 6) containing 0.75% NaCl or supplemented with 5%, 10%, and 30% NaCl at 4°C for 7 days were centrifugated at 10,000 × g for 3 min, rinsed three times with 0.1M Phosphate Buffered Saline (PBS, pH 7.2), and then re-suspended in 0.1M PBS to collect the pellets. These cells were fixed in 2% paraformaldehyde overnight at 4°C. After washing with PBS, cells of *V. parahaemolyticus* were post-fixed in 1% osmium tetroxide in 0.1M PBS and dehydrated in a series (30%, 50%, 70%, 95%, and 100%) of ethanol solutions. Thereafter, each of bacterial cells were infiltrated with 2 ml of epoxy resin. Polymerization of the resin was performed at 60°C for 24 hrs. Then, sections, approximately 120 nm thickness, were cut randomly and photographed with a JEOL JEM 1200 EX transmission electron microscope (JEOL USA Inc., Peabody, MA, USA).

### Fatty acid composition analysis

Whole cell fatty acid analysis was conducted according to a standardized method by Microbial Identification System (MIDI, Microbial ID Inc., Newark, Del., USA). Cells from 5 ml of a stationary-phase culture and 5 ml of VBNC *V. parahaemolyticus* were harvested by centrifugation at 10,000 × g for 5 min. Bacterial cells were subjected to a series of procedures, including saponification, methylation, and washing to extract carboxylic acid derivatives of long-chain aliphatic molecules. Then, extracted lipids were used to analyze the fatty acid composition of *V. parahaemolyticus* in the VBNC state, following several procedures provided by the supplier (Microbial Identification System, Newark, Del.)

### Assessment of VBNC bacterial resistances to an acidic environment

Ten ml of an exponential-phase culture and 10 ml of VBNC *V. parahaemolyticus* ATCC 17082were harvested by centrifugation at 10,000 × g for 5 min, washed twice, and then re-suspended in 10 ml of PBS, respectively. A simulated human gastric juice was prepared according a study conducted by Lee et al. (2016). This human digestion model (pH 1.5) consisted of 6.5 ml of HCl (37 g per 1 L of DW), 18 ml of CaCl_2_ · 2H_2_O (37 g per 1 L of DW), 1 g of bovine serum albumin (Corning), 2.5 g of pepsin (Sigma), and 3 g of mucin (Sigma). These chemicals and enzymes used in the present study were either autoclaved or filtered through a 0.2-ưm-size a polycarbonate membrane (ADVANTEC) before use. Thus, 10 ml of the bacterial cell was mixed with 10 ml of the gastric juice. Each of *V. parahaemolyticus* ATCC 17082 cells in a simulated stomach digestion model was incubated in a shaking water bath (120 rpm) at 37°C for 60 min. At a regular time-interval, the bacterial population was plating-counted on TSA and the viability of nonculturable *V. parahaemolyticus* ATCC 17082 was also enumerated with the fluorescence microscopic assay.

## ACKNOWLEDGEMENT

This research was supported by Basic Science Research Program through the National Research Foundation of Korea (NRF) funded by the Ministry of Science and ICT (NRF-2016R1A2B4014591).

## REFERENCES

1. Yue X, Liu B, Xiang J, Jia J. 2010. Identification and characterization of the pathogenic effect of a *Vibrio parahaemolyticus*-related bacterium isolated from clam *Meretrix meretrix* with mass mortality. J Invertebr Pathol 103:109–115.

2. Wong HC, Shen CT, Chang CN, Lee YS, Oliver JD. 2004. Biochemical and virulence characterization of viable but nonculturable cells of *Vibrio parahaemolyticus*. J Food Prot 67:2,430-2,435.

3. Yu WT, Jong KJ, Lin YR, Tsai S, Tey YH, Wong H. 2013. Prevalence of *Vibrio parahaemolyticus* in oyster and clam culturing environments in Taiwan. Int J Food Microbiol 160:185–192.

4. Hackney CR, Bay B, Speck ML. 1980. Incidence of *Vibrio parahaemolyticus* in and the microbiological quality of seafood in North Carolina. J Food Prot 43:769–773.

5. Piñeyro P, Zhou X, Orfe LH, Friel PJ, Lahmers K, Call DR. 2010. Development of two animal models to study the function of Vibrio parahaemolyticus Type III secretion systems. Infect Immun 78:4,551-4,559.

6. Yoon JH, Bae YM, Lee SY. 2017. Effects of varying concentrations of sodium chloride and acidic conditions on the behavior of *Vibrio parahaemolyticus* and *Vibrio vulnificus* cold-starved in artificial sea water microcosms. Food Sci Biotechnol 26:829–839.

7. Gribbon LT, Barer MR. 1995. Oxidative metabolism in nonculturable *Helicobacter pylori* and *Vibrio vulnificus* cells studies by substrate-enhanced tetrazolium reduction and digital image processing. Appl Environ Microbiol 61:3,379-3,384.

8. Mishra A, Taneja N, Sharma M. 2011. Demonstration of viable but nonculturable *Vibrio cholerae* O1 in fresh water environment of India using ciprofloxacin DFA-DVC method. Lett Appl Microbiol 53:124–126.

9. Liu Y, Wang C, Tyrrell G, Hrudey SE, Li XF. 2009. Induction of *Escherichia coli* O157:H7 into the viable but nonculturable state by chloraminated water and river water, and subsequent resuscitation. Environ Microbiol Rep 1:155–161.

10. Muela A, Seco C, Camafeita E, Arana I, Orruño M, López JA, Barcina I. 2008. Changes in *Escherichia coli* outer membrane subproteome under environmental conditions inducing the viable but nonculturable state. FEMS Microbiol Ecol 64:28–36.

11. Besnard V, Federighi M, Declerq E, Jugiau F, Cappelier J. 2002. Environmental and physico-chemical factors induce VBNC state in *Listeria monocytogenes*. Vet Res 33:359–370.

12. Cappelier JM, Minet J, Magras C, Colwell RR, Federighi M. 1999. Recovery in embryonated eggs of viable but nonculturable *Campylobacter jejuni* cells and maintenance of ability to adhere to HeLa cells after resuscitation. Appl Environ Microbiol 65:5,154-5,157.

13. Rahman I, Shahamat M, Chowdhury MA, Colwell RR. 1996. Potential virulence of viable but nonculturable *Shigella dysenteriae* type 1. Appl Environ Microbiol 62:115–120.

14. Chiang ML, Ho WL, Chou CC. 2008. Ethanol shock changes the fatty acid profile and survival behavior of *Vibrio parahaemolyticus* in various stress conditions. Food Microbiol 25:359–3765.

15. Rahman I, Shahamat M, Kirchman PA, Russek-Cohen E, Colwell RR. 1994. Methionine uptake and cytopathogenicity of viable but nonculturable *Shigella dysenteriae* type 1. Appl Environ Microbiol 60:3,573-3,578.

16. Antolinos V, Esteban MD, Ros-Chumillas M, Huertas JP, Periago PM, Palop A, Fernández PS. 2014. Assessment of the of acid shock effect on viability of *Bacillus cereus* and *Bacillus weihenstephanensis* using flow cytometry. Food Res Int 66:306–312.

17. Mizunoe Y, Wai SN, Ishikawa T, Takade A, Yoshida SI. 2000. Resuscitation of viable but nonculturable cells of *Vibrio parahaemolyticus* induced at low temperature under starvation. FEMS Microbiol Lett 115–120.

18. Dwidjosiswojo Z, Richard J, Moritz MM, Dopp E, Flemming HC, Wingender J. 2011. Influence of copper ions on the viability and cytotoxicity of *Pseudomonas aeruginosa* under conditions relevant to drinking water environments. Int J Hyg Environ Health 214:485–492.

19. Li N, Luo M, Fu YJ, Zu YG, Wang W, Zhang L, Yao LP, Zhao CJ, Sun Y. 2013. Effect of corilagin on membrane permeability of *Escherichia coli*, *Staphylococcus aureus*, and *Candida albicans*. Phytother Res 27:1,517-1,523.

20. Thennarasu S, Nagaraj R. 1996. Specific antimicrobial and hemolytic activities of 18-residue peptides derived from the amino terminal region of the toxin pardaxin. Protein Eng 9:1,219-1,224.

21. Zhang Z, Liu X, Wang Y, Jiang P, Quek SY. 2016. Antibacterial activity and mechanism of cinnamon essential oil against *Escherichia coli* and *Staphylococcus aureus*. Food Cont 59:282–289.

22. Balakumaran MD, Ramachandran R, Balashanmugam P, Mukeshkumar DJ, Kalaichelvan PT. 2016. Mycosynthesis of silver and gold nanoparticles: Optimization, characterization, and antimicrobial activity against human pathogens. Microbiol Res 182:8–20.

23. Linley E, Denyer SP, McDonnell G, Simons C, Maillard JY. 2012. Use of hydrogen peroxide as a biocide: new consideration of its mechanisms of biocidal action. J Antimicrob Chemother 67:1,589–1,596.

24. McDougald D, Gong L, Srinivasan S, Hild E, Thompson L, Takayama K, Rice SA, Kjelleberg S. 2002. Defences against oxidative stress during starvation in bacteria. Antonie Van Leeuwenhoek. 81:3–13.

25. Um HY, Kong HG, Lee HJ, Choi HK, Park EJ, Kim ST, Murugiyan S, Chung E, Kang KY, Lee SW. 2013. Altered gene expression and intracellular changes of the viable but nonculturable state in *Ralstonia solanacearum* by copper treatment. Plant Pathol J 29:374–385.

26. Abe A, Ohashi E, Ren H, Hayashi T, Endo H. 2007. Isolation and characterization of a cold-induced nonculturable suppression mutant of *Vibrio vulnificus*. Microbiol Res 162:130–138.

27. Nowakowska J, Oliver JD. 2013. Resistance to environmental stresses by *Vibrio vulnificus* in the viable but nonculturable state. FEMS Microbiol Ecol 84:213–222.

28. Klančnik A, Guzej B, Jamnik P, Vučković D, Abram M, Možina SS. 2009. Stress response and pathogenic potential of *Campylobacter jejuni* cells exposed to starvation. Res Microbiol 160:345–352.

29. Zhao F, Bi X, Hao Y, Liao X. 2013. Induction of viable but nonculturable *Escherichia coli* O157:H7 by high pressure CO_2_ and its characteristics. PLOS One 8:e6238.

30. Jia J, Chen J, Jiang Y, Tang J, Yang L, Liang C, Jia Z, Zhao L. 2014. Visualized analysis of cellular fatty acid profiles of *Vibrio parahaemolyticus* strains under cold stress. FEMS Microbiol Lett. 357:92–98.

31. Day AP, Oliver JD. 2004. Changes in membrane fatty acid composition during entry of *Vibrio vulnificus* into the viable but nonculturable state. J Microbiol. 42:69–73.

32. Beattie SH, Williams AG. 1999. Detection of toxigenic strains of *Bacillus cereus* and other *Bacillus* spp. with an improved cytotoxicity assay. Lett Appl Microbiol 28:221–225.

33. Jeßberger N, Dietrich R, Bock S, Didier A, Märtlbauer E. 2014. *Bacillus cereus* enterotoxins act as major virulence factors and exhibit distinct cytotoxicity to different human cell lines. Toxicon 77:49–57.

34. Dektas E, Daferera D, Sökmen M, Serdar G, Ertürk M, Polissiou MG, Sökmen A. 2016. In vitro antimicrobial, antioxidant, and antiviral activities of the essential oil and various extracts from *Thymus nummularis* M. Bieb. Indian J Tradit Knowl 15:403–410.

35. Wong HC, Wang P, Chen SY, Chiu SW. 2004. Resuscitation of viable but non-culturable *Vibrio parahaemolyticus* in a minimum salt medium. FEMS Microbiol Lett 233:269–275.

